# Neurocomputational mechanisms underlying fear-biased adaptation learning in changing environments

**DOI:** 10.1101/2022.06.25.497438

**Authors:** Zhihao Wang, Tian Nan, Katharina S. Goerlich, Yiman Li, André Aleman, Yuejia Luo, Pengfei Xu

**Author notes:** Corresponding authors: Pengfei Xu, Ph.D., Faculty of Psychology, Beijing Normal University, No.19, Xinjiekouwai St, Haidian District, Beijing 100875, China.; Yuejia Luo, Ph.D., Shenzhen Key Laboratory of Affective and Social Neuroscience, Shenzhen University, No.3688 Nanhai Ave., Nanshan District, Shenzhen 518060, China. The authors declare no competing financial interests.

## Abstract

Humans are able to adapt to the fast-changing world by estimating statistical regularities of the environment. Although fear can profoundly impact adaptive behaviors, the neural mechanisms underlying this phenomenon remain elusive. Here, we conducted a behavioral experiment (n = 21) and a functional magnetic resonance imaging experiment (n = 37) with a novel cue-biased adaptation learning task, during which we simultaneously manipulated emotional valence (fearful/neutral expressions of the cue) and environmental volatility (frequent/infrequent reversals of reward probabilities). Across two experiments, computational modelling consistently revealed a higher learning rate for the environment with frequent versus infrequent reversals following neutral cues. In contrast, this flexible adjustment was absent in the environment with fearful cues, suggesting a suppressive role of fear in adaptation to environmental volatility. This suppressive effect was underpinned by activity of the posterior parietal cortex, ventral striatum, hippocampus and dorsal anterior cingulate cortex (dACC) as well as increased functional connectivity between the dACC and temporal-parietal junction (TPJ) for fear with environmental volatility. Dynamic causal modelling identified that the driving effect was located in the TPJ and was associated with dACC activation, suggesting that the suppression of fear on adaptive behaviors occurs at the early stage of bottom-up processing. These findings provide a neuro-computational account of how fear interferes with adaptation to volatility during dynamic environments.

## Introduction

Humans and animals are able to adapt to the fast-changing world. Fear, the most studied emotion despite differences regarding its definition and measurement^1–4^ profoundly influences adaptive behavior^5,6^. Although adaptation to dynamic environments is a key form of behavioral flexibility, how fear affects adaptation to dynamic environments remains unclear. Evolutionarily, fear acts as an alerting signal for self-protection^7^. A rich literature illustrates beneficial effects of fear on flexibility^2,6,8,9^. Thus, elicitation of fearful signals may facilitate adaptation to dynamic environments. In contrast, when dominating consciousness, fear can disrupt systems supporting flexible behavior^5,10^, and lesions of fear circuits may even facilitate flexible performance^11^. Consequently, the conscious experience of fear may suppress adaptation to changing environments.

Adaptation to dynamic environments depends on the internal representation of uncertainties^12^, which can be classified into two types: expected uncertainty and unexpected uncertainty^12–14^. The former refers to noise in the action-outcome association, for example when choosing the correct option occasionally results in an undesirable outcome. The latter is characterized as volatility, or the frequency at which action-outcome contingencies change. For instance, after the switch of the action-outcome association, an action that was primarily associated with a given outcome becomes predominantly associated with another. Optimal adaptive behavior depends on accurate identification of the source of uncertainty^14–17^. More specifically, if unexpected outcomes are caused by noise, the current action is optimally guided by averaging previous observations. Instead, if unexpected outcomes result from environmental volatility, only recent outcomes are necessary to determine the present action. According to reinforcement learning theory^18^, human learners can adapt to changing environments, exhibiting a higher learning rate in the volatile relative to stable environment^14,16^. The Bayesian learner has also been demonstrated to dynamically track environmental volatility with optimal performance^14,17^. Significant steps forward in uncertainty-related studies point toward the relevance of affective representations^19–21^. Emotional responses contribute to adaptation to volatility^14^, and failure to adapt to environmental volatility has been identified as a major contributor to affective disorders^17,22–24^. Animal studies demonstrated a facilitating role of serotonin neurons in flexible adaptation to volatility^19^ and observed that levels of the neurotransmitter serotonin modulated fear processing^25^, suggesting an important link between fear and adaptation to volatility.

Neuroimaging studies have identified a set of brain areas linking emotion to adaptation to environmental volatility. The dorsal anterior cingulate cortex (dACC) represents subjective estimation of environmental volatility^14,26^. In addition to a critical role in uncertainty computation^13,27–29^, signals from the dACC have also been linked to affective responses and in particular to integrative processing from multiple sources (i.e., emotion-cognition integration)^30–32^. The amygdala has been shown to encode associations under uncertain environments, especially for Pavlovian conditioning^33^. As a fear circuitry hub, the amygdala has been shown to mediate interactions between fearful reactions and executive functions^30^. The hippocampus (HI) has long been considered crucial for fear memory^6,34^. Learning signals during uncertainty are evidenced to be sorted in the HI, highlighting an important role of the HI in adaptation to changing environments^13,35^. The ventral striatum (VS), a key region for reward processing, is broadly implicated in affective learning and value representation^6,36,37^. The VS has also been identified to adjust human flexible responses when the aversive environment changed^33^. The orbitofrontal cortex (OFC) shows high sensitivity to the processing of uncertainty and plays an important role in emotion-related subjective value representation and updating in both human and animals^38–40^.

This study aimed to examine the effect of fear on adaptation to volatility and its underlying neuro-computational mechanisms. From an evolutionary perspective, it can be hypothesized that fear would facilitate adaptation to volatility learning. Alternatively, fear could prevent adaptation to volatility learning due to the cognitive cost of fear processing and a flight response induced by fearful stimuli with which the individual wants to exit the volatile environment. To test these hypotheses, we used computational modelling and functional magnetic resonance imaging (fMRI) in a novel cue-biased adaptation learning task. Based on the framework of the probabilistic reward reversal learning task^14,24^, the current task simultaneously manipulated emotional valence of the cue (fearful/neutral facial expressions) and environmental volatility (frequent/infrequent reversals of reward probability)^16^. Please note that the presentation of facial expressions in a brief period of time to participants is the classic emotional induction method^41,42^. We observed a higher learning rate for the environment with frequent compared to infrequent reversals with the cue of neutral face, which was consistent with previous studies^14,17,24^. However, this pattern was absent in the fearful cues, suggesting a suppressive role of fear in adaptation to volatility and thus supporting the second hypothesis. We also revealed the distributed neural substrates underlying computations of fear-biased adaptation to volatility learning, including integration systems, learning systems, and memory systems. In particular, the TPJ-dACC pathway was found to mediate the interplay between fear and adaptation to volatility learning. This may imply that the suppression of fear may affect adaptive behaviors at a relatively early stage of processing bottom-up inputs.

## Results

Participants completed the cue-biased adaptation learning task (Figure 1) in Experiment 1 (exp1; n=21, behavioral study) and Experiment 2 (exp2; n=40, fMRI study; see Table 1 for demographic information). Using the framework of the probabilistic reward reversal learning task^14,24^, we developed a two [emotional valence of cue: fearful/neutral expressions (fear/neut)] by two [environmental volatility: frequent/infrequent reversals (freq/infreq); Figure 1] within-subject design to test how fear influences adaptation to volatility.

**Figure1.**
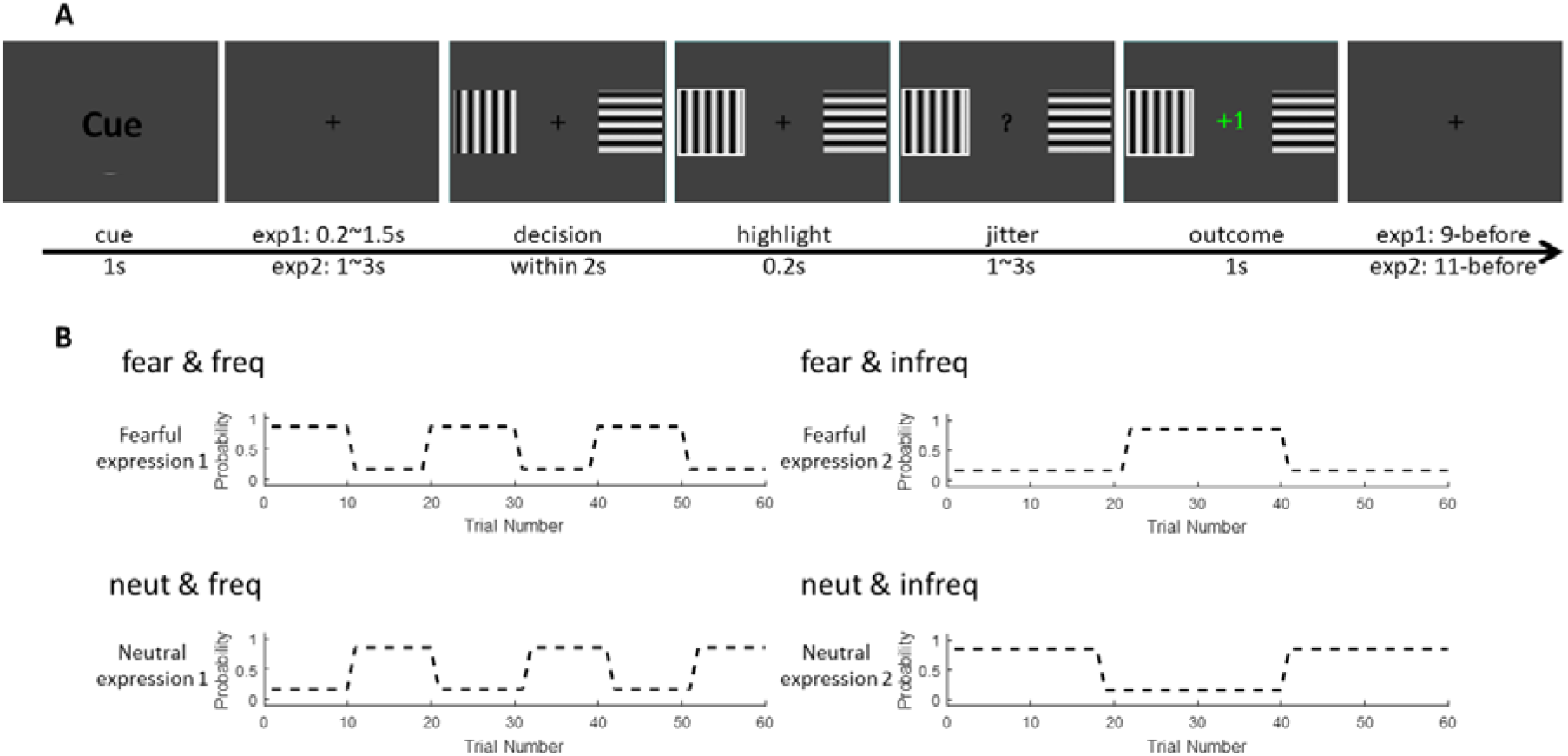
Experimental design. Trial design of the fear-biased volatility learning task (A) and an example of contingency between cue and environmental volatility (B).

**Table 1.** Demographics and questionnaire scores

|  | exp 1 (n = 21) | exp 2 (n = 40) |
| --- | --- | --- |
| Age | 20.81 (1.94) | 21.75 (2.13) |
| Gender<br>(female) | 11 | 18 |
| BVAQ | 122.71 (8.76) | 124.70 (10.71) |
| BVAQ-cog | 74.90 (7.87) | 74.10 (7.28) |
| BVAQ-aff | 47.81 (6.31) | 50.60 (5.69) |
| MASQ-AA | 21.95 (6.90) | 23.03 (5.91) |
| MASQ-AD | 63.10 (14.05) | 59.90 (12.33) |
| MASQ-GDA | 15.14 (5.63) | 16.68 (5.04) |
| MASQ-GDD | 18.14 (6.22) | 18.73 (7.34) |

Descriptive data are presented as mean (SD). Abbreviations: BVAQ, the Bermond-Vorst Alexithymia Questionnaire; BVAQ-cog, the cognitive dimension of BVAQ; BVAQ-aff, the affective dimension of BVAQ; MASQ-AA, anxious arousal of the Mood and Anxiety Symptoms Questionnaire; MASQ-AD, anhedonia depression; GDA, general distress: anxiety; GDD, general distress: depression.

### Manipulation checks

We first tested the recognition rate of fearful/neutral expressions of the cue. We collected post-ratings of “To which extent do you think the presented face showed a fearful/neutral expression?” using a Likert scale from 0 (totally not) to 8 (totally agree). One-sample *t*-tests confirmed that post-ratings were significantly higher than the random level (i.e., 4: midpoint between 0 and 8) both for fearful and neutral expressions (exp1: *ts* > 9.608, *ps* < 0.001; exp2: *ts* > 8.308, *ps* < 0.001; Figure S1; Table S2), suggesting the participants correctly identified the emotional expressions of the cue. To examine whether participants were able to learn the reward structure of our task, in line with Piray et al. (2019)^43^, we plotted participants’ performance after reversal. Results showed that participants indeed learned the reward schedule successfully (Figure 2 A and B). In addition, a small number of missing trials (less than 14) and a low ratio of short response time (< 200 ms; < 1.67%; see Table S2) suggested sufficient engagement during the task.

**Figure2.**
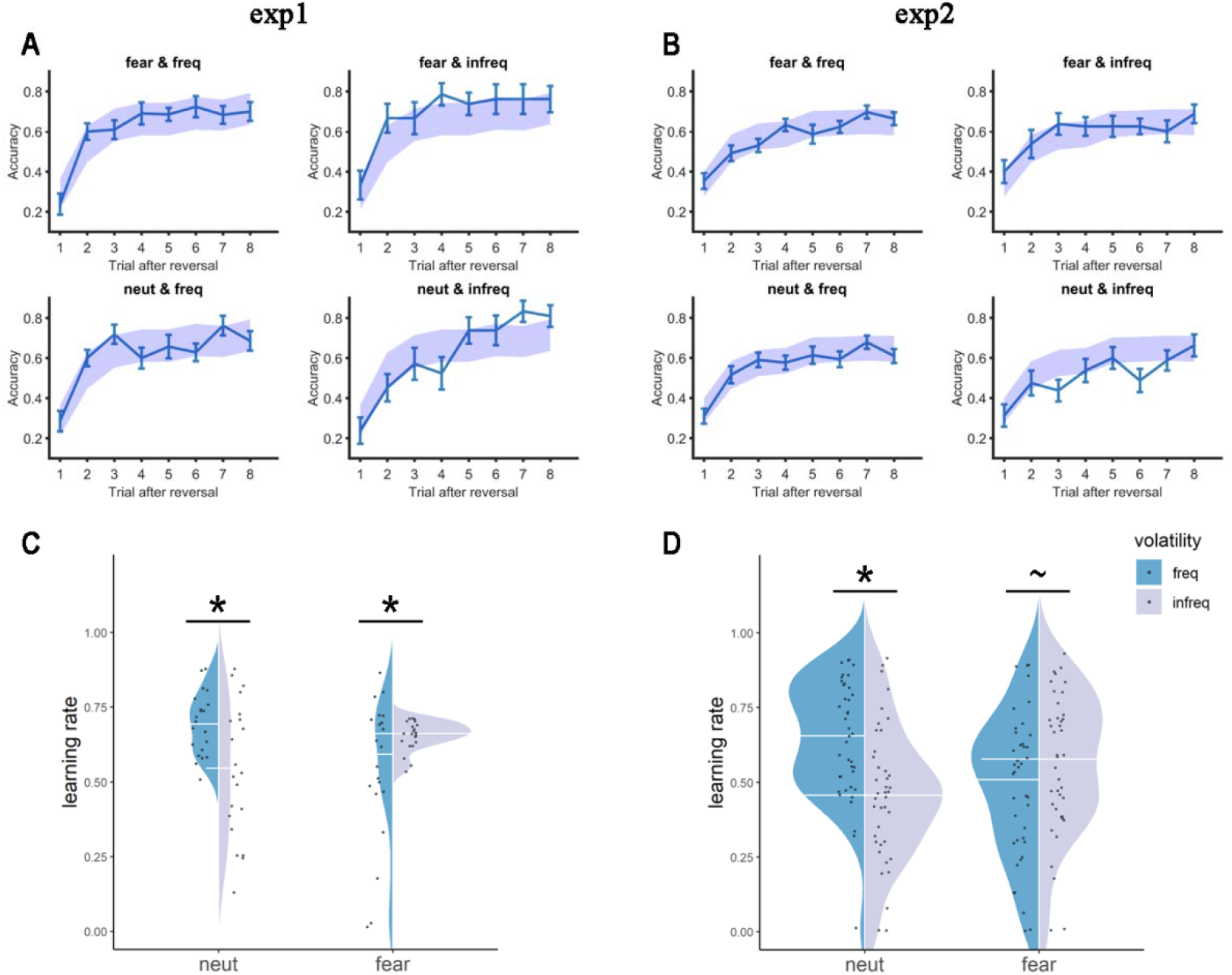
Behavioral results. (AB) Performance after reversals. Solid lines represent real data (mean ± standard error). Shaded panels represent 95% highest density interval (HDI) from the winning model. (CD) Learning rate from the winning model. Note: \**p* < 0.05; ^~^*p* < 0.1.

### Computational mechanisms underlying fear-biased adaptation to volatility learning

To formally quantify cognitive mechanisms underlying fear-biased adaptation to volatility learning, we constructed 11 models, with considerations of learning by trial and error (Rescorla-Wagner model), attentional lapses, forgetting, and learning by attention (Pearce-Hall model; see Methods and Materials). Using indices of leave-one-out information criterion (LOOIC) and widely applicable information criterion (WAIC), model comparison showed that the winning model was Model 1 (M1) in both exp1 and exp2 [except slightly better for model 5 (M5) at the index of LOOIC in exp2, ΔLOOIC = −0.4; Table 2]. Specifically, M1 assumed that each condition was learned differently with the trial and error strategy (one learning rate per condition). Model simulation from M1 showed high correlation coefficients between real accuracy and simulated accuracy for every condition in both exp1 and exp2 (exp1: *rs* > 0.82, *ps* < 0.001; exp2: *rs* > 0.85, *ps* < 0.001; Figure S2). Simulated data closely resembled the real performance (Figure 2A and B), confirming the validity of parameter estimation.

**Table 2.** Model comparison. The winning model in both exp1 and exp2 is M1.

| Models | Number of parameters | exp 1 (n = 21) |  | exp 2 (n = 40) |  |
| --- | --- | --- | --- | --- | --- |
| | | ΔLOOIC | ΔWAIC | ΔLOOIC | ΔWAIC |
| M1 | 8 | 0 | 0 | 0 | 0 |
| M2 | 4 | 11.3 | 20.7 | 103.4 | 118.5 |
| M3 | 5 | 7.6 | 23.0 | 35.9 | 53.2 |
| M4 | 8 | 64.9 | 68.2 | 112.2 | 126.8 |
| M5 | 9 | 24.6 | 57.8 | -0.4 | 11.5 |
| M6 | 9 | 118.7 | 89.7 | 107.6 | 112.5 |
| M7 | 10 | 74.5 | 85.1 | 136.4 | 135.2 |
| M8 | 10 | 62.0 | 70.4 | 123.7 | 128.1 |
| M9 | 7 | 53.6 | 63.0 | 124.9 | 131.6 |
| M10 | 8 | 53.9 | 59.0 | 123.9 | 131.0 |
| M11 | 8 | 51.7 | 58.1 | 112.8 | 124.4 |

Abbreviations: ΔLOOIC, leave-one-out information criterion relative to the winning
model; $\Delta\text{WAIC}$ , widely applicable information criterion relative to the winning model.

To examine how learning rates from the winning model M1 were modulated by emotional cue and environmental volatility, we implemented linear mixed-effect models (LMM) with subject as a random factor and with cue (fear/neut) and volatility (freq/infreq) as within-subject factors for exp1 and exp2, respectively. Results consistently showed increased learning rates in freq as compared to infreq in neutral cues (exp1: F_1,20_ = 15.433, *p* = 0.001, partial *η^2^* = 0.436, Figure 2C; exp2: F_1,39_ = 21.711, *p* < 0.001, partial *η^2^* = 0.358; Figure 2D), replicating previous results regarding adaptation to volatility learning^14,17,24^. However, this pattern disappeared in the face of fearful cues (exp1: F_1,20_ = 5.460, *p* = 0.030, partial *η^2^* = 0.214; freq < infreq; Figure 2C; exp2: F_1,39_ = 3.501, *p* = 0.069, partial *η^2^* = 0.082; Figure 2D; Table S3). These findings suggest that fear interferes with adaptation to volatility learning. In addition, M5, adding a component of attentional lapse, performed slightly better than M1 at the index of LOOIC in exp2 (ΔLOOIC = −0.4; Table 2). The implementation of LMM for M5 showed the same pattern as M1 (Figure S4), suggesting that the current results were highly robust. Note that statistic values for main effects and interaction effects are reported in the Supplementary Materials.

### Neural correlates of fear-biased adaptation to volatility learning: Learning rate

To examine how fear-biased adaptation to volatility learning was represented in the human brain from the perspective of learning rate, we performed the first generalized linear model (GLM1). The behavioral bias was defined as [(fear & freq – fear & infreq) – (neut & freq – neut & infreq)] in learning rates, whereas the neural bias was defined as [(fear & freq – fear & infreq) – (neut & freq – neut & infreq)] in BOLD signals at the outcome stage. This study focused on the ventral striatum (VS), dorsal anterior cingulate cortex (dACC), amygdala, orbitofrontal cortex (OFC), and hippocampus (HI) that have been identified to link emotion to adaptation to environmental volatility (see Introduction for details). These brain regions were thus hypothesized to contribute to fear-biased adaptation learning. ROI analysis showed positive correlations between the behavioral bias and the neural bias in the VS (ROI analysis, peak at [4 8 −4], r = 0.578, *p* < 0.001, k = 7; Figure 3B), and HI (ROI analysis; peak at [28 −16 −22]; r = 0.606, *p* < 0.001, k = 10; Figure 3C). No significant activation in the amygdala, dACC and OFC was found. We also conducted exploratory whole-brain analysis, revealing a positive correlation between the behavioral bias and the neural bias in the posterior parietal cortex (PPC; whole-brain analysis; peak at [−50 −60 50] in MNI coordinates; r = 0.583, *p* < 0.001, k = 103; Figure 3A). These results suggest that the PPC, VS, and HI encode fear-biased adaptation to volatility learning.

**Figure3.**
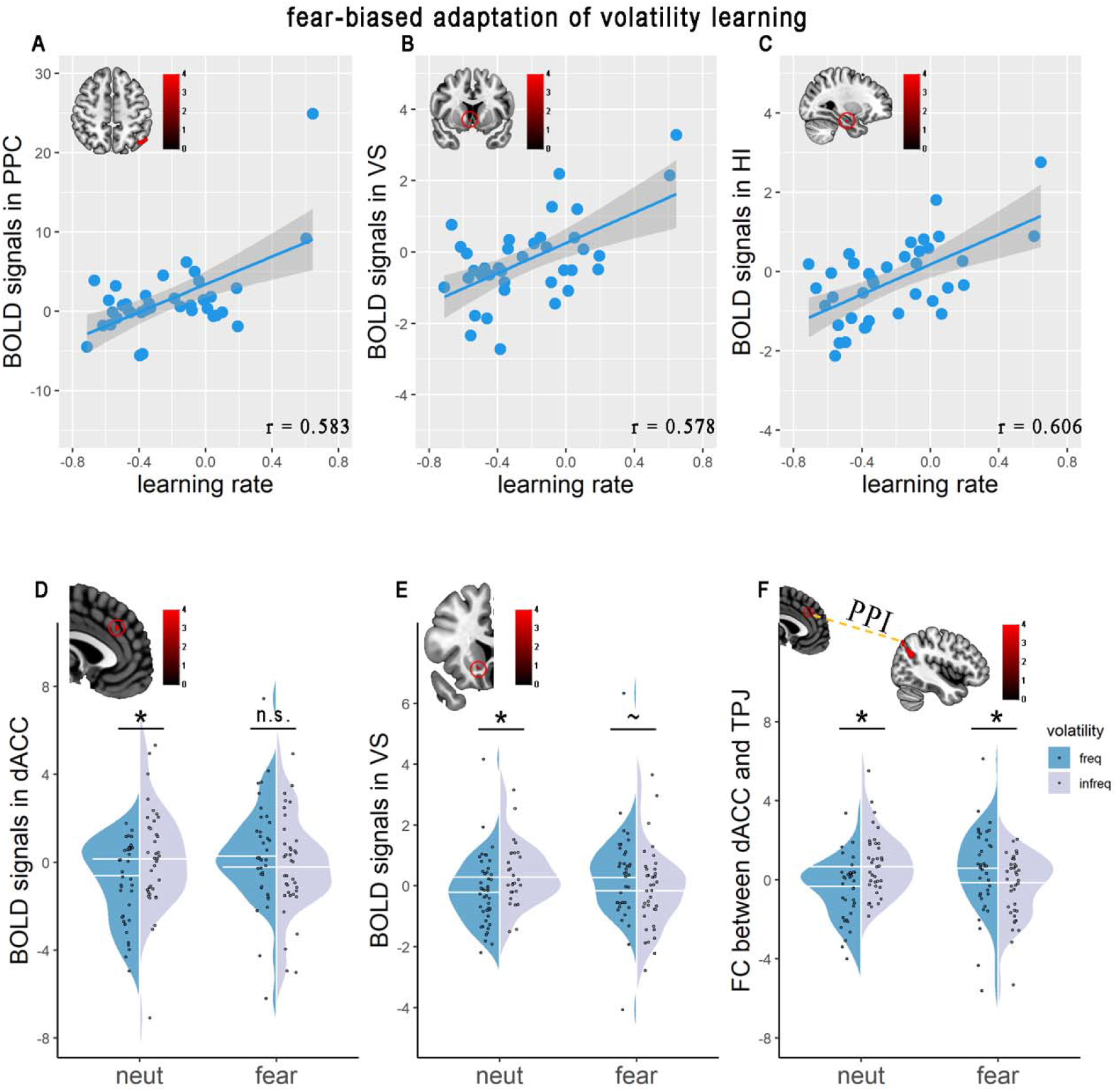
Activity and connectivity results. (ABC) Neural correlates of fear-biased adaptation to volatility learning in terms of learning rate in PPC (peak at [−50 −60 50], k = 103, A), VS (peak at [−4 8 −4], k = 7, B), and HI (peak at [28 −16 −22], k = 10, C). (DE) activation results from fear-biased adaptation to volatility learning in terms of subjective volatility in dACC (peak at [−2 32 32], k = 8, D) and VS (peak at [10 16 −8], k = 9, E). (F) functional connectivity between dACC (seed region) and TPJ (target region, peak at [−64 −52 36], k = 263) for each condition. Circles in red represent region-of-interest (ROI) analysis. Note: PPC, posterior parietal cortex; VS, ventral striatum; HI, hippocampus; dACC, dorsal anterior cingulate cortex; n.s., not significant; *\*p* < 0.05; ^~^*p* < 0.1.

### Neural substrates of fear-biased adaptation to volatility learning: Subjective volatility

To examine representations of the human brain for fear-biased adaptation to volatility learning, we implemented model-based fMRI analysis (GLM2) with parametric modulation of subjective volatility at the stage of outcome. Again, the neural bias was defined as [(fear & freq – fear & infreq) – (neut & freq – neut & infreq)] in the parametric activation of subjective volatility. Note that trial-by-trial subjective volatility was derived from Bayesian Learner model (see Methods and Materials for details; see Figure S3 for an example participant). We observed significant activations in the dACC (ROI analysis; peak at [−2 32 32]; k = 8, Figure 3D) and VS (ROI analysis; peak at [10 16 −8]; k = 9; Figure 3E). No significant activation in the amygdala, HI and OFC was found. The implementation of ANOVAs in these significant activations checked the direction of interaction effects. For both dACC and VS, we observed significant increases in infreq versus freq in neutral cues (dACC: F_1,36_ = 13.272, *p* = 0.001, partial *η^2^* = 0.269; Figure 3D; VS: F_1,36_ = 8.191, *p* = 0.007, partial *η^2^* = 0.185; Figure 3E). However, such patterns disappeared for fearful cues (dACC: F_1,36_ = 1.833, *p* = 0.184, partial *η^2^* = 0.048; Figure 3D; VS: F_1,36_ = 3.966, *p* = 0.054, partial *η^2^* = 0.099; Figure 3E). These findings indicate that the dACC and VS are engaged to encode subjective volatility in fear-biased adaptation.

We next sought to explore the underlying brain networks modulating fear-biased adaptation to volatility learning. Based on model-based fMRI results, generalized psychophysiological interaction (gPPI)^44^ analysis was performed with the dACC and VS as seed regions, separately. We found significant connectivity of the dACC with the temporal parietal junction (TPJ; whole-brain analysis; peak at [−64 −52 36]; k = 263; Figure 3F). The interaction effect further showed that functional connectivity between dACC and TPJ increased in infreq as compared to freq following neutral cues (F_1,36_ = 11.306, *p* = 0.002, partial *η^2^* = 0.239; Figure 3F), but decreased following fearful cues (F_1,36_ = 4.748, *p* = 0.036, partial *η^2^* = 0.117; Figure 3F). The gPPI results suggest that the dACC-TPJ circuit encodes subjective volatility calculations in fear-biased adaptation.

To further test directions of information flow between the dACC and TPJ underlying fear-biased adaptation to subjective volatility, we conducted dynamic causal modelling (DCM) analysis^45^. We constructed six models with different assumptions of modulatory effects and driving effects while we fixed the full intrinsic connectivity (Figure 4A; for Methods for details). The random-effect (RFX) Bayesian Model Selection (BMS) using indices of exceedance probability and expected posterior probability recommended model 3 as the winning model (Figure 4B&C). The model 3 assumed a driving effect from the TPJ, a modulatory effect from the TPJ to dACC, and another modulatory effect from the dACC to TPJ. We found a significant interaction effect between cue and environmental volatility in the driving effect on the TPJ (F_1,36_ = 8.321, *p* = 0.007, partial η^2^ = 0.188; Figure 4D). Simple effect analysis showed a significant increase in infreq than freq in neutral cues (F_1,36_ = 9.827, *p* = 0.003, partial *η^2^* = 0.214; Figure 4D), but not fearful cues (F_1,36_ = 0.064, partial *η^2^* = 0.002; Figure 4D), suggesting the modulation of the driving effect from the TPJ on fear-biased adaptation to subjective volatility. Furthermore, the driving bias from the TPJ was positively correlated to parametric bias for BOLD signals in subjective volatility in the dACC (r = 0.371, *p* = 0.024; Figure 4E). Taken together, consistent with our behavioral modelling results, these neural results relating to subjective volatility support the notion that fear interferes with adaptation to volatility learning and further uncover the driven computation from the TPJ in the dACC-TPJ pathway underlying fear-biased adaptation to volatility learning.

**Figure4.**
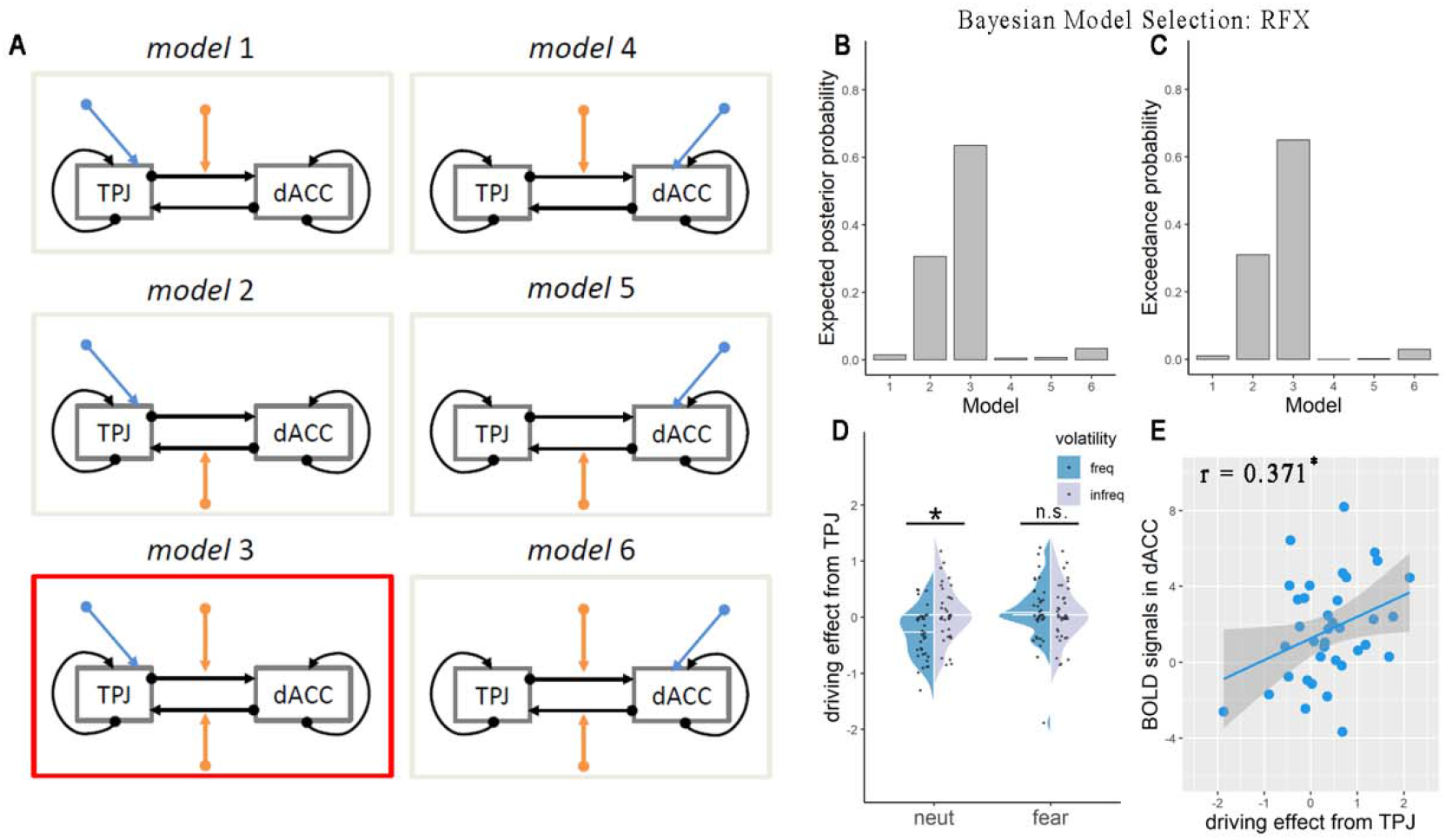
DCM results. (A) Model space for DCM analysis. Black arrows represent intrinsic connectivity. Blue arrows represent the driving effect. Orange arrows represent modulatory effects. The winning model was highlighted in red. Model comparison using indices of expected posterior probability (B) and exceedance probability (C). (D) Driving effect on TPJ for each condition. (F) Correlation between driving effect on TPJ and parametric activation of subjectivity volatility in dACC in terms of fear-biased adaptation to volatility learning. Note: TPJ, temporal parietal junction; dACC, dorsal anterior cingulate cortex; RFX, random-effect; n.s., not significant; \**p* < 0.05.

### Both cognitive and affective alexithymia were related to fear-biased adaptation to volatility learning

We also took into account individual differences in propensity for emotion processing and regulation, as this may be a factor of interest. More specifically, this regards alexithymia, which refers to a reduced ability to identify, describe and regulate one’s feelings^46^. Given cognitive and emotional deficits in alexithymia^47^, we explored associations between alexithymia levels and fear-biased adaptation to volatility learning. We found a significant correlation between the total score of the Bermond-Vorst Alexithymia Questionnaire (BVAQ)^48^ and the behavioral bias (in learning rate; r = 0.351, *p* = 0.006; Figure 5A). Please note that higher scores on the BVAQ represented lower levels of alexithymia. The higher score for behavioral bias [(fear & freq – fear & infreq) – (neut & freq – neut & infreq)] reflected weaker influence of fear on adaptation to volatility learning. Therefore, the positive correlation suggests that the interference of fear with adaptation to volatility learning was stronger the more alexithymic individuals were. To examine contributions of cognitive (BVAQ-cog) and affective dimensions (BVAQ-aff) of alexithymia for learning rate bias, we performed correlations of learning rate bias with BVAQ-cog and BVAQ-aff, respectively. We observed (marginally) significant correlations for BVAQ-cog (r = 0.291, *p* = 0.023; Figure 5B) and BVAQ-aff (r = 0.227, *p* = 0.078; Figure 5C), indicating that both cognitive and affective dimensions of alexithymia explained fear-biased adaptation to volatility learning. In addition, anxiety and depression has been demonstrated to co-occur with alexithymia. We, therefore, performed partial correlations to control for the potential influences of anxiety and depression. Correlation coefficients remained significant for total BVAQ scores, affective dimension scores and cognitive scores (*rs* > 0.269, *ps* < 0.043). In sum, individuals prone to either cognitive or affective alexithymia were more influenced by fear in adaptation to volatility.

**Figure5.**
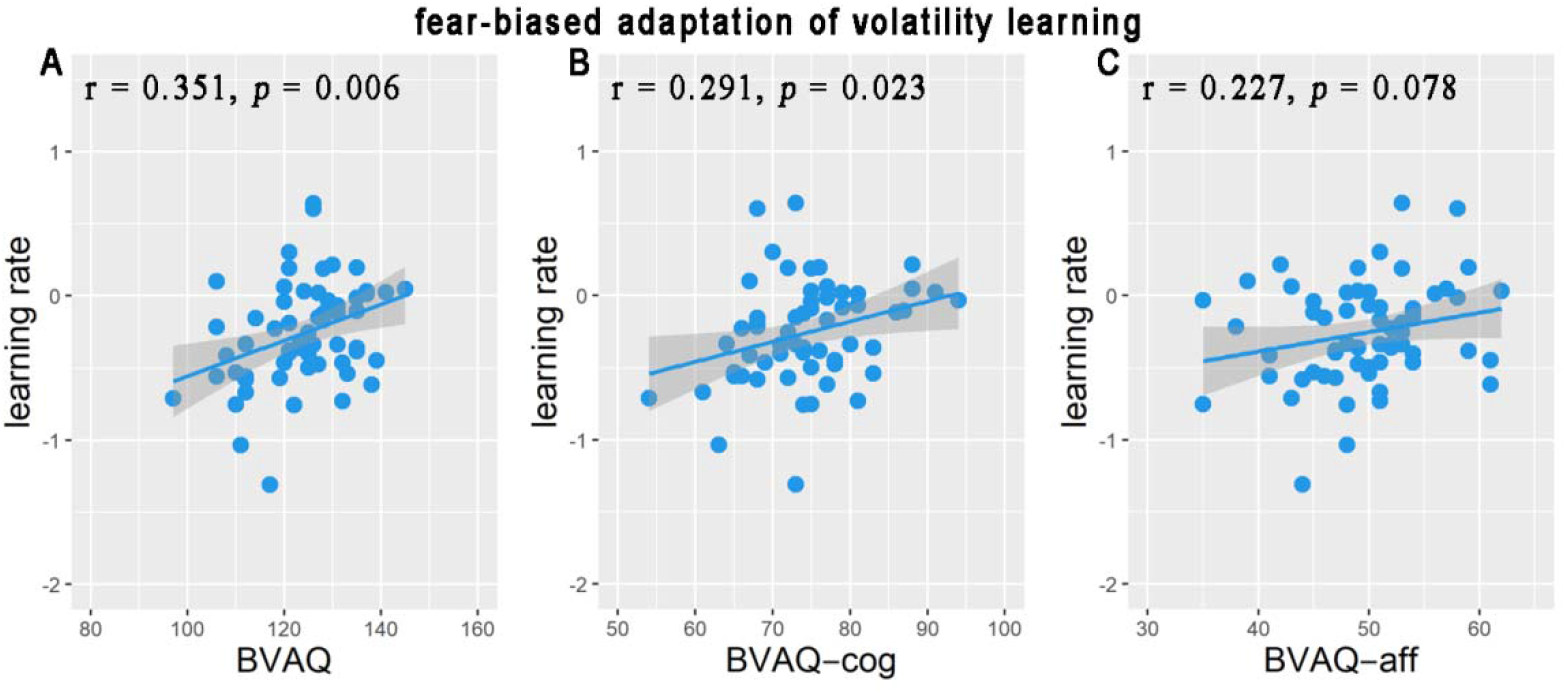
Correlations of learning rate in terms of fear-biased adaptation to volatility learning with BVAQ (A), BVAQ-cog (B), and BVAQ-aff (C). Note: BVAQ, the Bermond-Vorst Alexithymia Questionnaire; BVAQ-cog, cognitive dimension of BVAQ; BVAQ-aff, affective dimension of BVAQ.

## Discussion

This study examined neuro-computational mechanisms of how fear influences adaptation to volatility. In two independent experiments, we consistently found that the environment with frequent reversals elicited a higher learning rate than the environment with infrequent reversals in neutral cues, replicating previous studies^14,17,24^. However, this difference was absent in the face of fearful cues, supporting the hypothesis that fear prevented adaptation to volatility. This suppressive effect was underpinned by activity of the posterior parietal cortex, ventral striatum, hippocampus and dorsal anterior cingulate cortex (dACC), as well as functional connectivity between the dACC and temporal-parietal junction (TPJ), suggesting distributed brain systems for computations of fear-biased adaptation. Effective connectivity results further showed that this bias mainly resulted from the driving effect of the TPJ, indicating that computations underlying fear-biased adaptation to volatility may occur at a relatively early stage of processing bottom-up inputs. Lastly, this bias was stronger with higher levels of alexithymia.

Using computational modelling, this study revealed that fear inhibits adaptation to volatility. In our control conditions of neutral cues, participants showed a higher learning rate to the environment with frequent (as compared to infrequent) reversals, consistent with previous findings of a higher learning rate for volatile than stable conditions^14,17,24^. This finding supports the notion that humans are able to adjust learning rates in adaptation to volatility^14,16,17,24^. Critically, this adjustment was disrupted by fearful signals. It has been shown that fear disrupts systems supporting flexible behavior through dominating consciousness^6,10,11^. Therefore, the conscious experience of fear potentially inhibits adaptive function, in particular adaption to volatility. This supports the notion of mutual inhibition between emotion and cognition and the dual competition model of emotion-cognition integration^30,49,50^. However, some studies showed that emotion, especially fear, promoted cognitive flexibility^6,8,9^. The contextual factor likely contributes to this discrepancy. Specifically, our findings of the suppressive effect of fear on adaptation to volatility were in the rewarding situation. It is widely believed that fear facilitates performance during the threatening environment^2,6,7^. There is evidence that fear is preferentially activated in aversive contexts^2,7^. Considering the design of our study, the current findings may be limited to the rewarding context. Future studies would benefit from investigating the interplay between fear and adaptation to volatility in threatening contexts.

Neuroimaging results showed that the dACC encodes fear-biased adaptation to volatility learning. In our control conditions, decreased activation in the dACC was observed in environments with frequent versus infrequent reversals. This finding was contrary to previous findings of a positive correlation of dACC signals with subjective volatility^14,26^. It has been proposed that task difficulty mediates the relationship between activation in the dACC and cognitive demand in an inverted U-shape pattern^42,51^. One explanation for the discrepancy in the activation of the dACC might be that the current task was more difficult than the previous probabilistic reward learning task due to the design of interleaved trials from four conditions. More importantly, the signal of volatility in the dACC was absent in fearful conditions, suggesting a suppressive role of fear in the adaptive function of the dACC. The dACC is considered to be a higher-order integrative hub for multiple information^30–32^. For example, integration between fearful signals and executive functions has been found in the dACC^30^. Therefore, these results indicate that fear modulates the dACC’s signals to track environmental volatility and support the integrative processing of emotion and cognition in the dACC.

The TPJ was also found to work in concert with the dACC to mediate fear-biased adaptation to volatility in our study. A nexus model of the TPJ has contended a basic integrative hub for the TPJ function, including attentional processing, memory storage, and social processing^52^. For instance, the TPJ was identified to be involved in global Gestalt integration at the perceptive level^53^. Moderation of dACC-TPJ functional connectivity on fear-biased adaptation to volatility in our study suggests a circuit of information interchange between low- and high-level integrative processing of fear and adaptation learning. DCM analysis further showed bidirectional information flow between the dACC and TPJ during fear-biased adaptation learning. Importantly, fear-biased adaptation to volatility was modulated by the driving effect from the TPJ. This may suggest that fear mainly impedes bottom-up information input and low-level integrative processing between fear and adaptation to volatility. In sum, in combination with the significant correlation between the activated bias in the dACC and the driving bias in the TPJ (Figure 5E), these results reveal a crucial role of the TPJ-dACC neural pathway in volatility learning of fear-biased adaptation.

We also observed activation in the VS, but not the amygdala, in fear-biased adaptation to volatility, which may suggest differential roles of the VS and amygdala. Both the VS and amygdala have been demonstrated to represent uncertainties during dynamic environments and emotion processing^33,36,37,54^. Within learning contexts, the amygdala is linked to Pavlovian conditioning, while the VS has been implicated in instrumental learning^6^. The current results may thus reflect the interference of instrumental adaptation by fear in the VS. Another explanation regards a possible role of Pavlovian-Instrumental Transfer. This transfer effect underlying the amygdala and VS has been demonstrated in animal models^6,55^. In this study, Pavlovian tendencies were gradually dominated by instrumental learning. This transfer effect thus explains activation in the VS, but not the amygdala, in fear-biased adaptation learning. However, interpretations about the negative result of activation in the amygdala should be cautious. In addition, the HI and PPC were found to moderate fear-biased adaptation to volatility in our study. The HI has been thought to encode and store memory^56,57^, while the PPC was implicated in memory retrieval^58^. These two brain regions were also involved in learning and value representation in the uncertain environment^13,35,59,60^. Therefore, these results may indicate that fear interferes with memory signals in the HI and PPC for adaptation to dynamic environments. Our study did not find any significant OFC activation. This is surprising given its crucial involvements in emotion-related valuation and flexible learning^38–40^. One potential explanation is low signal-noise ratio for the OFC BOLD signals^61^. In brief, distributed brain areas involved in memory and learning systems seem to contribute to fear-biased adaptation to volatility learning.

Unexpectedly, no direct correlation between learning rate and subjective volatility was observed in this study. The Bayesian Learner model has been proposed in a hierarchical structure^14^ with i) subjective estimation of environmental volatility at the 1st level, ii) subjective estimation of the winning probability at the 2nd level, and iii) the observed outcome at the 3rd level. Based on the reinforcement learning model without hierarchical processing, learning rates were fit from the subjective probability across trials at the 2nd level. Previous studies indeed showed a positive correlation between volatility signals and learning rate^14,17^. However, a recent simulation study demonstrated that learning rates were jointly estimated from both stochasticity and volatility^16^. Although separate correlations of activation in the HI were observed with the volatility index and the learning rate, the absence of a direct correlation between volatility and learning rate may be due to the ignorance of stochasticity in our model space, which was in line with previous studies^14,17,24^. Future studies may benefit from taking stochasticity into account when constructing computational models to examine the relationship between learning rate and volatility.

We observed that maladaptation to volatility following fear cues correlated with alexithymia, suggesting a stronger influence of fear on adaptation to volatility learning in individuals with higher levels of alexithymia. Alexithymia refers to difficulties in emotion processing^46^. Alexithymia has been considered as a transdiagnostic risk factor for various affective disorders, such as anxiety and depression^62,63^. While a rich literature showed both cognitive and emotional deficits in alexithymia^47,42,64,65^, the current results further suggest an integrative difficulty between fear and adaptation to volatility with higher alexithymia levels in healthy individuals. Given that abnormalities in regulating fearful signals are at the core of affective disorders^6^ and failure to appropriately adjust learning to dynamic environments contributes to affective disorders^17,22–24^, our findings provide additional insights into pathological mechanisms of alexithymia-related affective disorders. However, in our recent study, we did not find impaired emotion-cognition integration in alexithymia^66^. One possible explanation may be that the stimuli used in our previous study merely displayed isolated parts of the human body and thus lacked social-emotional features^67^, whose processing seems to be particularly impaired in relation to alexithymia^65^. Therefore, impaired fear-biased adaptation to volatility in alexithymia may be social-specific.

Several limitations of the present study should be mentioned. First, we elicited different emotions only using social stimuli, e.g., facial expressions. This study complied with Marr’s 3-level theory at the computation, algorithm, and implementation levels^68^, addressing the question of how fear influences adaptation to volatility. However, given that social specificity is at the core of Marr’s framework, whether our findings of the neuro-computational mechanisms underlying fear-biased adaptation to volatility were social-specific remains unclear. In addition, three types of fear may be distinguished^2^: 1) fear of physical objects, e.g., thunder; 2) fear of other humans; 3) fear of animals, e.g., spider. These types of fear are corresponding to nature phobias, social phobias, and animal phobias, respectively^2^. Our findings only provide insights into social fear. Next, activity of the locus coeruleus norepinephrine system cannot be detected with the current device and parameters. This system has been implicated in tracking environmental volatility and generating emotional reactions^17,69^. Future studies would benefit from exploring the function of this system in fear-biased adaptation to volatility learning using eye-tracking techniques.

To conclude, this study provides a neuro-cognitive account of how fear interferes with adaptation to volatility during dynamic environments at the computational level. We also show that this bias is related to individual differences in propensity for emotion processing and regulation. Our findings reveal distributed brain substrates underlying fear-biased adaptation to volatility, including integration systems, learning systems, and memory systems. The TPJ-dACC pathway, in particular, mediates the interplay between fear and adaptation to volatility learning, suggesting that the suppression of fear on adaptive behaviors may occur at a relatively early stage of processing bottom-up inputs. Our work thus sheds light on the neuro-computational mechanisms underlying fear-biased adaptation to volatility learning.

## Materials and Methods

### Participants

A total of 64 right-handed healthy adults at Shenzhen University participated in two experiments. Experiment 1 (exp1) was a behavioral study, including 21 participants (11 females, age = 20.81±1.94). Experiment 2 (exp2) was an fMRI study, including 43 participants. Due to impatience (one participant), and many missing trials (two participants failed to respond in more than 10% of the trials; 62 and 46 in the 240 trials, respectively), and excessive head motion (three participants with more than 10% scans with frame-wise displacement (FD) > 0.5)^70^, the final sample for exp2 included 40 participants (18 females, age = 21.75±2.13) for behavioral analysis and 37 participants (16 females, age = 21.57±2.06) for fMRI analysis. See Table 1 for demographic information. All the participants had no history of neurological and psychiatric disorders or head injury. The study was approved by the local Ethics Committee at Shenzhen University and written informed consent was obtained from all participants.

### General Procedure

After signing the informed consent, participants were asked to complete a three-stage training task to understand the probability and reversal components underlying the task^71^. The cue-biased adaptation learning task was then implemented (while participants in exp2 simultaneously experienced fMRI scanning). After the task, participants were asked to rate each fearful and neutral expression by answering the question “To which extent do you think the presented face showed a fearful/neutral expression?”. These post-ratings with a Likert scale from 0 (totally not) to 8 (totally agree) were used to check whether participants correctly recognized these stimuli as fearful/neutral expressions. Next, participants completed several questionnaires, including the Chinese version of the Bermond-Vorst Alexithymia Questionnaire (BVAQ)^48^ and the Mood and Anxiety Symptoms Questionnaire (MASQ)^72^. Finally, participants were paid based on their learning performance, which was instructed before the experiment.

### Cue-biased adaptation learning task

Inspired by Behrens et al. (2007)^14^ and Piray et al. (2019)^43^, we developed a cue-biased adaptation learning task (Figure 1). Using the framework of the probabilistic reward reversal learning task^14,24^, we manipulated the type of cue (fearful/neutral expressions) and environmental volatility (frequent/infrequent reversals)^16^. Specifically, frequent reversals (freq) were defined as a situation in which the contingency reversed every 9~11 trials randomly, while infrequent reversals (infreq) referred to a situation in which the contingency reversed every 18~22 trials randomly. In brief, the current study with a two by two within-subject design consisted of 240 trials, with 60 trials per condition. Please note that trials for each condition were intermixed within a block. Participants were asked to learn by trial and error to maximize their payoff. Although participants were explicitly informed that reward structure would change throughout the task, they needed to infer the moment on which reversal occurred and the speed at which reversal changed^73^. At the beginning of each trial, a cue (a fearful or neutral face) was first shown on the screen center with duration of 1 s (Figure 1A). To eliminate potential gender effects, facial expressions with the same gender with participants were used as cues. For example, the female face with the fearful or neutral expression was shown on the cue for the female participants. We selected four female faces and four male faces from the Taiwanese Facial Expression Image Database (TFEID)^74^, with two fearful and neutral expressions for each gender. The normative rating (category and intensity) from the TFEID can be seen in Table S1. Please note that the cue type and the environmental volatility were randomly matched across participants. After a fixation cross with a random duration (0.2~1.5 s for exp1 and 1~3 s for exp2), two options (horizon and vertical Gabor patches) were presented randomly on each side. Participants were required to make a decision within 2 s. Upon response, the selected option would be highlighted for 0.2 s, followed by a question mark at the screen center with a jitter interval at 1~3 s. Then the outcome was shown for 1 s, with “+1” in green indicating reward and “+0” in red reflecting no reward. Outcome was delivered with a probability of reward at either 85% or 15%, based on the outcome schedule (Figure 1B). If participants did not respond within 2 s, which was defined as a missing trial, the option would not be highlighted and outcome was “+0”. At the end of a trial, a fixation cross was presented for 0.3 ~ 5.6 s to ensure that each trial lasted for 9 s for exp1 and 0.8 ~ 6.8 s to ensure that each trial lasted for 11 s for exp2. Exp1 and exp2 lasted for 36 and 44 minutes, respectively. All experimental procedures were presented using E-prime 2.0 (Psychology Software Tools Inc. Pittsburgh, PA, USA).

### Self-report Questionnaires

To assess alexithymia, we used the Chinese version of the BVAQ^48^. This questionnaire consists of 35 items, each answer being scored on a five-point Likert scale of 1 (this in no way applies to me) to 5 (this definitely applies to me). BVAQ includes both cognitive and affective components with acceptable reliability and validity. Higher scores for BVAQ represented lower levels of alexithymia. To control potential confounding effects of anxiety and depression, participants also completed the Chinese version of the Mood and Anxiety Symptoms Questionnaire (MASQ)^72^. The MASQ consists of 62 items that are assessed with a four-point Likert scale of 1 (not at all) to 4 (extremely). It measures symptoms of anxiety and depression based on a tripartite model, including general distress, which can be further divided into general distress: anxiety (GDA) and general distress: depression (GDD), anxious arousal (AA), and anhedonia depression (AD) subscales.

### Behavioral data analysis

For validation, one-sample *t* tests were performed in post-ratings for fearful and neutral expression, separately. We also plotted performance after reversal to check whether participants learned the reward structure of our task, in line with Piray et al. (2019)^43^.

### Computational modeling of task performance

To best describe participants’ learning performance in our fear-biased volatility learning task, we conducted a stage-wise model construction procedure^24,75^. That is, we added each component to the model or modified an existing component progressively, based on the best model from the previous stage. Model comparison used the leave-one-out information criterion (LOOIC) and the widely applicable information criterion (WAIC) to avoid overfitting. Lower scores of LOOIC or WAIC indicated better out-of-sample prediction accuracy of the candidate model. Parameter estimation was performed using hierarchical Bayesian analysis. Posterior inference was performed with Markov chain Monte Carlo (MCMC) sampling with 4000 iterations across 4 chains from the posterior distribution. The entire modelling-related procedures were performed using the hBayesDM package^76^. In total, we tested 11 candidate models using the stage-by-stage model construction procedure in exp1. For exp2, we compared these models directly.

We used the simple Rescorla-Wagner (RW) model (Equation 1-3) as the baseline model^18^, which was widely used in learning-related studies^14,17,24^. The simple RW model assumed that participants learned reward structure by trial and error (Equation 1-3)^18^.

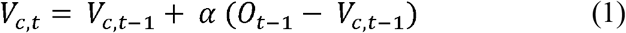

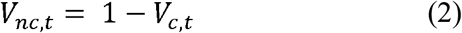

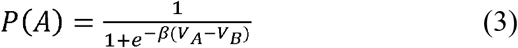

Here, the value V of the chosen option is updated trial-by-trial, which are determined by both the prediction error and learning rate *α* (0 < *α* < 1). The prediction error is derived from the difference between received outcome (O) and expected value (V) from the previous trial. For simplicity, the value of the unchosen option is regarded as the opponent value for the chosen option^14,17^. Finally, we used a softmax function with a decision parameter *β* (or inverse temperature parameter; 0 < *β* < 10) to calculate the chosen probability for each option.

In Stage 1, we compared a model (M1) that assumed each condition was learned differently to a model (M2) that assumed that participants learned differently due to volatility^14^, but not the type of cue. That is, M2 assumed that there was no cue (emotional) effect for all parameters. Thus, M1 included a learning parameter (learning rate) and a decision parameter (inverse temperature) for each condition, whereas M2 consisted of 2 learning parameters and decision parameters for conditions of frequent and infrequent reversals. Given that it was well established that the environment with frequent reversals elicited higher learning rate than that with infrequent reversals^14,17,24^, we thus did not include the model assuming that there was no effect of volatility. M1 with eight parameters showed better performance than M2 with four parameters (see Table 2).

In Stage 2, we removed/added some components to the M1 to validate the winning model from Stage 1. M3 assumed that participants learned differently for each condition but shared a decision parameter across conditions (five parameters). M4 assumed that participants regarded our task as the two-arm bandit and only updated the value of the chosen option (eight parameters), rather than the one-arm bandit (Equation 2). M5 added a parameter of the lapse in attention (*ε*; 0 < *ε* < 1; Equation 4), assuming that participants occasionally made random choices due to a lapse in attention (nine parameters).

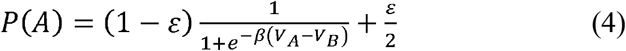

Model comparisons among M1, M3, M4, and M5 indicated that M1 performed better (see Table 2).

Participants needed to learn four types of cue-option-outcome contingencies in our task (as compared to one type in the previous probabilistic reward reversal learning task), including the option of forgetting^77^. Therefore, in Stage 3, we added forgetting parameters *ϕ* to the M1 (0 < *ϕ* < 1; Equation 5). This parameter pulled the estimated value toward the random level (0.5).

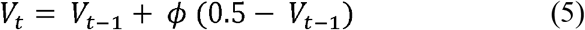

M6 assumed a sharing forgetting parameter across conditions (nine parameters). M7 assumed *ϕ* were modulated by volatility, with 2 forgetting parameters for environments with frequent and infrequent reversals (ten parameters). Rather, we assumed *ϕ* were modulated by emotional cues in M8 (ten parameters). Model comparisons among M1, M6, M7, and M8 showed that M1 performed better (see Table 2).

In Stage 4, we considered hybrid models of the Pearce-Hall model with the RW model given that this type of hybrid model performed best among the candidate models during the emotion- and volatility-related task^43^. M9 assumed a shared weighting parameter ω across conditions (0 < *ω* < 1; seven parameters). M10 assumed distinct weighting parameters *ω* for fearful and neutral cues (eight parameters). M11 assumed different scale parameters *κ* of learning rate for fearful and neutral conditions (0 < *κ* < 1; 8 parameters). M9-11 were the same as M2, M4, and M5 in Piray et al. (2019), respectively. Again, M1 won among these candidate models (see Table 2).

We further performed model validation for the winning model (M1) in exp1 using the generated data from MCMC sampling (4000 iterations). First, we computed correlations between real accuracy and simulated accuracy for each condition. Please note that mean simulated accuracy across 4000 iterations per condition and participant was used. Second, we analyzed the generated data using the index of performance after reversal and computed 95% high density posterior interval (HDI) to compare the difference between simulated data and real data.

Next, we compared these 11 candidate models directly in exp2. The same procedures of model validation were performed for exp2 behavioral data.

### Statistical analysis for learning rate

Each parameter was represented by the mean of the posterior distribution of the parameters. We conducted LMMs on learning rate with subject as a random factor and with cue (fear/neut) and volatility (infreq/freq) as within-subject factors for exp1 and exp2, respectively. Pearson correlations of fear-biased adaptation to volatility learning in terms of learning rate were implemented with total BVAQ scores, the cognitive dimension, and the affective dimension of BVAQ scores across 61 participants. Statistical analyses were conducted using SPSS 17.0 (IBM Inc). We set the significance level at *p* = 0.05.

### Bayesian Learner model

Bayesian Learner model has been shown to dynamically track environmental volatility^14^. Specifically, the estimated probability for the option can be inferred from the observed outcome (reward or not; Equation 6).

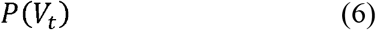

Please note that *V_t_* can also be represented as probability given that the magnitude in our study was fixed at 1. The probability for the current trial could be inferred from the probability of the previous trial and perceived volatility of the current trial (Equation 7).

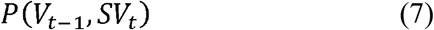

where *SV_t_* is the subjective volatility at the *t^th^* trial.

Subjective volatility of the current trial can be inferred from those of the previous trial and a distrust variable *K_t_* in the current trial (Equation 8).

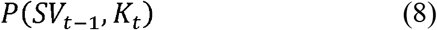

Consistent with previous studies^14,17^ and for theoretic considerations, we also derived trial-by-trial subjective volatility from the Bayesian Learner model using analytic inference^14^.

### Image acquisition and Preprocessing

MRI data were acquired with a Siemens Trio 3T scanner at Shenzhen. Both the fMRI and high-resolution 3D structural brain data were obtained using a 64-channel phased-array head. The fMRI data were acquired by means of a gradient-echo echo-planar imaging sequence containing the following parameters: repetition time (TR) = 1000 ms, echo time (TE) = 30 ms, 78 multiband slices, slice thickness 2 mm with gap 2 mm, flip angle = 35°, field of view (FOV) = 192 mm × 192 mm, data matrix = 96 × 96, and 240 volumes scanned in 2640 seconds. Additionally, the 3D structural brain images (1mm^3^ isotropic) were acquired for each participant using a T1-weighted 3D magnetization-prepared rapid gradient echo sequence with the following parameters: TR / TE = 2300 ms / 2.26 ms, flip angle = 8°, data matrix = 232 × 256, FOV = 232 mm × 256 mm, BandWidth = 200 Hz / pixel, 192 image slices along the sagittal orientation, obtained in about 9 min. Functional MRI data were preprocessed with DPABI^78^ (http://rfmri.org/dpabi), a software package based on SPM12 (version no.7219; https://www.fil.ion.ucl.ac.uk/spm/software/spm12/). It comprised the following steps: 1) realignment; 2) co-registering the T1-weighted image to the corresponding mean functional image; 3) segmenting into grey matter, white matter and cerebrospinal fluid by DARTEL; 4) normalizing to the standard Montreal Neurological Institute space (MNI template, resampling voxel size 2×2×2 mm^3^); 5) smoothing with a Gaussian kernel of 6 mm full width at half maximum (FWHM).

### Generalized Linear Models (GLM)

SPM12 was used for general linear model (GLM) analysis. The current study focused on the interaction effect between cue (fear/neut) and volatility (infreq/freq), or fear-biased adaptation to volatility learning, which was defined as [(fear & freq – fear & infreq) – (neut & freq – neut & infreq)].

#### Neural responses to learning rates

Based on the winning model and its parameter estimation, the learning rate for each condition (mean value across 4000 iterations) was obtained. The first model (GLM1) aimed to examine fear-biased adaptation to volatility learning at the neural level in terms of learning rates. The first-level design matrix in GLM1 included the cue for fearful and neutral expressions, option presentation, highlight of the selected options, jitter, and outcome for each condition. If any, we modeled missing trials with option presentation, highlight of the selected options, jitter, and outcome. In addition, the six head motion parameters and a FD regressor were included as covariates of no interest. The regressors were then convolved with the canonical hemodynamic response function (HRF). To obtain the fear-biased adaptation to volatility learning effect, we performed one-sample *t*-tests for contrasted images [(fear & freq – fear & infreq) – (neut & freq – neut & infreq)] at outcome onsets. These contrasted betas further regressed against fear-biased adaptation to volatility learning in terms of learning rate [(fear & freq – fear & infreq) – (neut & freq – neut & infreq)] at the second level.

#### Neural responses to subjective volatility

Trial-by-trial subjective volatility from Bayesian Learner model was used to examine neural correlates of fear-biased adaptation to volatility learning in terms of subjective volatility (GLM2). The first-level design matrix in GLM2 included the cue for fearful and neutral expressions, option presentation with the parametric modulation by subjective volatility, highlight of the selected options, jitter with the parametric modulation by subjective volatility, and outcome with the parametric modulation of subjective volatility for each condition. If any, we modeled missing trials with option presentation, highlight, jitter, and outcome. In addition, the six head motion parameters and a FD regressor were included as covariates of no interest. The regressors were then convolved with the canonical HRF. Finally, contrasted images [(fear & freq - fear & infreq) - (neut & freq - neut & infreq)] at outcome onsets for the modulator effect of subjective volatility from the first-level analysis were entered into one-sample *t*-tests for the second-level analysis.

### Generalized Psychophysiological Interaction (gPPI)

To assess how the functional connectivity between BOLD signals in the seed region and BOLD signals in the target region was modulated by fear-biased adaptation to volatility learning, we performed generalized psychophysiological interaction (gPPI) using the gPPI toolbox^44^. The seed region(s) were determined by activation results from GLM2. Specifically, we extracted the deconvolved (with HRF) time course from the first eigenvariate of the seed region (the physiological term) for each participant. The subjective volatility at outcome onset for each condition was used as the psychological term. The interaction term was then generated by multiplying the physiological term with the psychological term. The convolved interaction term was entered into the GLM analysis. All imaging results were corrected with the threshold of *p* < 0.001 at the voxel level and with the threshold of *p* < 0.05 at the cluster level using AlphaSim procedure. The dACC was selected from Behrens et al. (2007)^14^ (MNI: x = −6, y = 26, z = 34 mm, a sphere with 10mm radium), whereas other ROIs were obtained from AAL atlas. Small volume correction (SVC) was used for regions of interest (ROIs).

### Dynamic Causal Modelling (DCM)

To further assess effective connectivity between the brain regions underlying the modulation of fear-biased adaptation, we performed dynamic causal modelling (DCM) using the DCM12 toolbox^45^. We used the same parametric signals (subjective volatility) from each condition to test fear-biased adaptation effects. Six models were constructed with different assumptions of modulatory effects (B matrix) and driving effects (C matrix) while we fixed the full intrinsic connectivity (A matrix; Figure 4A). For the winning model, ANOVAs were performed with cue (fear/neut) and volatility (freq/infreq) as within-subject factors in A, B and C matrices, respectively. Bonferroni correction was used to correct for multiple comparisons.

## Acknowledgements

This study was funded by the National Natural Science Foundation of China (31920103009, 31871137, 32071083, 32020103008, and 32071100), the Major Project of National Social Science Foundation (20&ZD153), Young Elite Scientists Sponsorship Program by China Association for Science and Technology (YESS20180158), Natural Science Foundation of Guangdong Province (2020A1515011394), Shenzhen-Hong Kong Institute of Brain Science – Shenzhen Fundamental Research Institutions (2019SHIBS0003), and Shenzhen Science and Technology Research Funding Program (JCYJ20180507183500566 and JCYJ20190808121415365).

## Data availability

The data that support the findings of this study are available from the corresponding author upon reasonable request.

