## Supplement for "Neurocomputational mechanisms underlying fear-biased adaptation learning in changing environments"

**Statistic values for main effects and interaction effects**

Regarding learning rates in exp1, we observed a significant interaction effect between cue and environmental volatility (F_1,20_ = 19.095, *p* < 0.001, partial $\eta^{2}$ = 0.24). No significant main effects were found (main effect of cue: F_1,20_ = 0.349, partial $\eta^{2}$ < 0.001; main effect of environmental volatility: F_1,20_ = 0.569, partial $\eta^{2}$ < 0.001). With regard to learning rates in exp2, we observed a significant interaction effect between cue and environmental volatility (F_1,39_ = 22.509, *p* < 0.001, partial $\eta^{2}$ = 0.16). No significant main effect of cue were found (F_1,39_ = 1.130, *p* = 0.290, partial $\eta^{2}$ < 0.001) in addition to the main effect of environmental volatility (F_1,39_ = 3.954, *p* = 0.049, partial $\eta^{2}$ = 0.03). In terms of parametric effects of subjective volatility in the dACC, we observed a significant interaction effect between cue and environmental volatility (F_1,36_ = 14.609, *p* < 0.001, partial $\eta^{2}$ = 0.289). No significant main effects were found (main effect of cue: F_1,36_ = 1.763, *p* = 0.193, partial $\eta^{2}$ = 0.047; main effect of environmental volatility: F_1,36_ = 0.933, partial $\eta^{2}$ = 0.025). As for parametric effects of subjective volatility in the VS, we observed a significant interaction effect between cue and environmental volatility (F_1,36_ = 15.274, *p* < 0.001, partial $\eta^{2}$ = 0.298). No significant main effects were found (main effect of cue: F_1,36_ = 0.021, partial $\eta^{2}$ = 0.001; main effect of environmental volatility: F_1,36_ = 0.090, partial $\eta^{2}$ = 0.002). For functional connectivity between the dACC and TPJ, we observed a significant interaction effect between cue and environmental volatility (F_1,36_ = 26.551, *p* < 0.001, partial $\eta^{2}$ = 0.424). No significant main effects were found (main effect of cue: F_1,36_ = 0.172, partial $\eta^{2}$ = 0.005; main effect of environmental volatility: F_1,36_ = 0.455, partial $\eta^{2}$ = 0.012). Regarding the driving effect on the TPJ, we found a significant main effect of cue (F_1,36_ = 8.321, *p* = 0.007, partial $\eta^{2}$ = 0.188; fear > neut) and a significant interaction effect between cue and environmental volatility (F_1,36_ = 8.321, *p* = 0.007, partial $\eta^{2}$ = 0.188; Figure 4D). No significant main effect of environmental volatility was observed (F_1,36_ = 2.476, *p* = 0.124, partial $\eta^{2}$ = 0.064).

**Figure S1.** Post-ratings for fearful and neutral expressions. Note: **p* < 0.05.

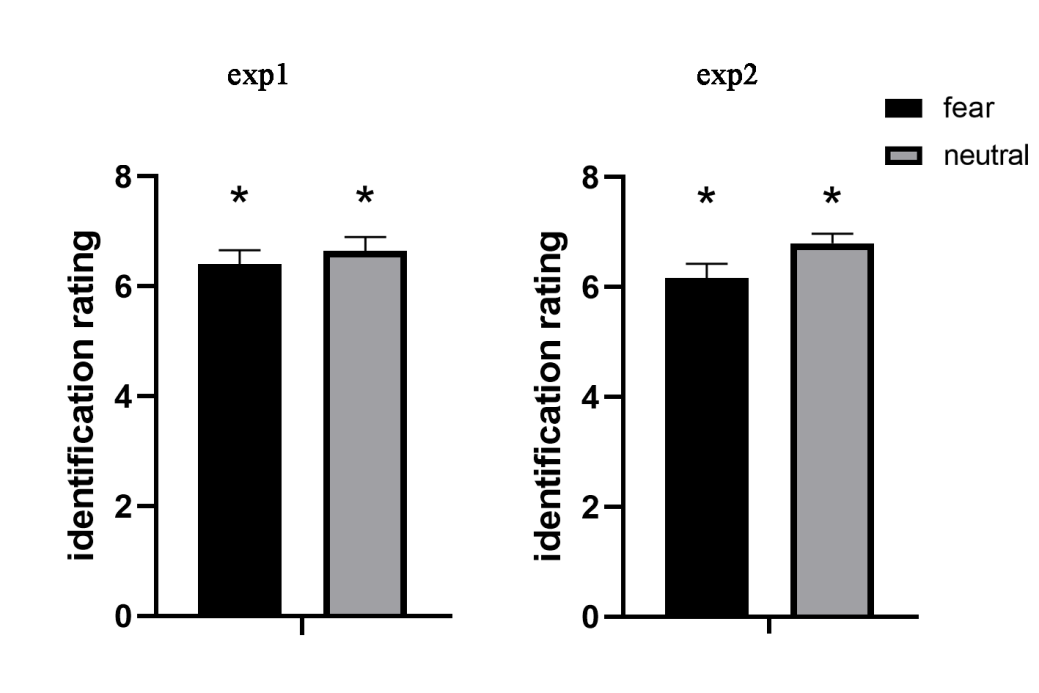

**Figure S2.** Model validation using the correlation between real accuracy and simulated accuracy.

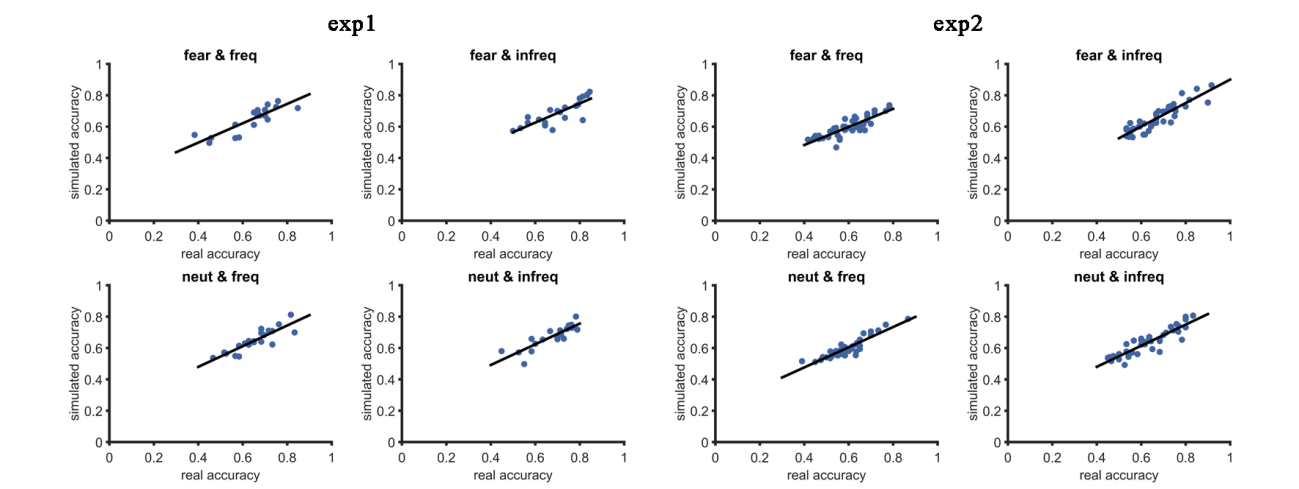

**Figure S3.** Subjective volatility estimated from Bayesian Learner model in an example participant. Solid lines in blue represent trial-by-trial estimated volatility. Dash lines represent reward schedules.

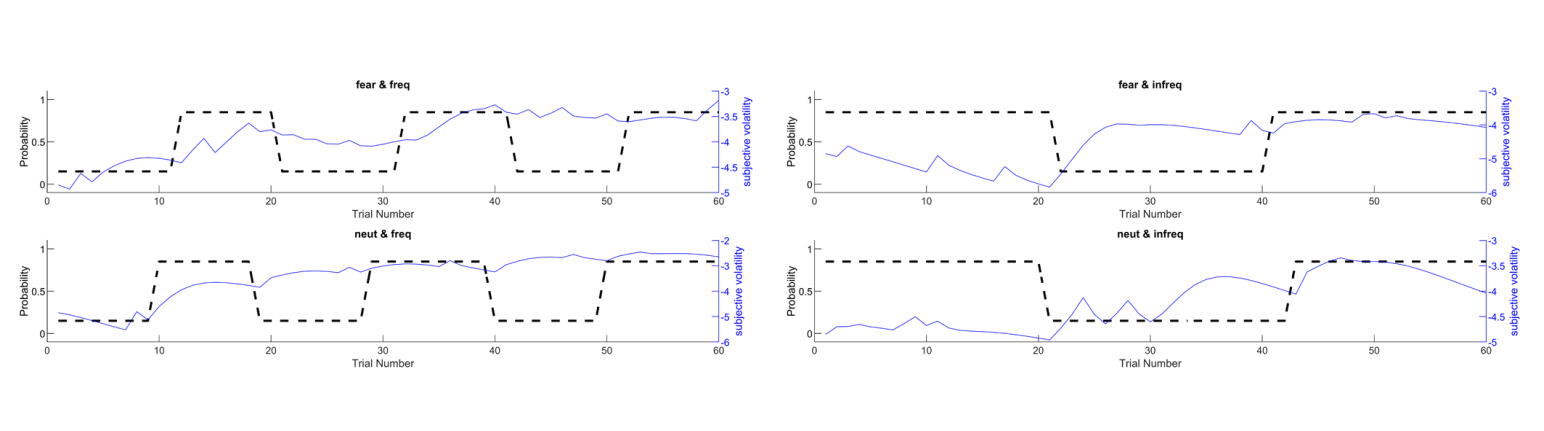

**Figure S4.** Learning rate from M5 in exp2. Note: n.s., not significant; **p* < 0.05.

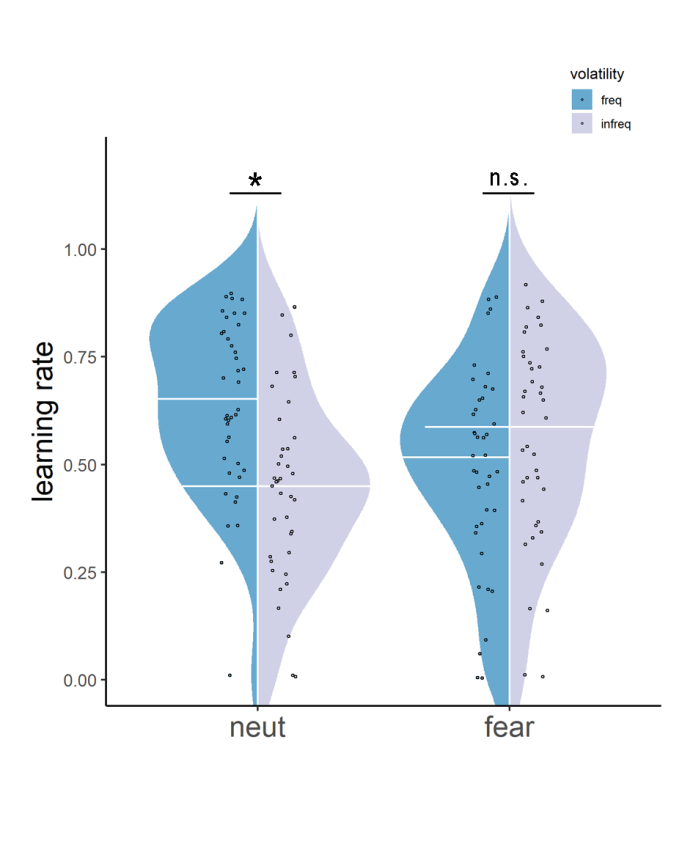

**Figure S5.** Other correlations.

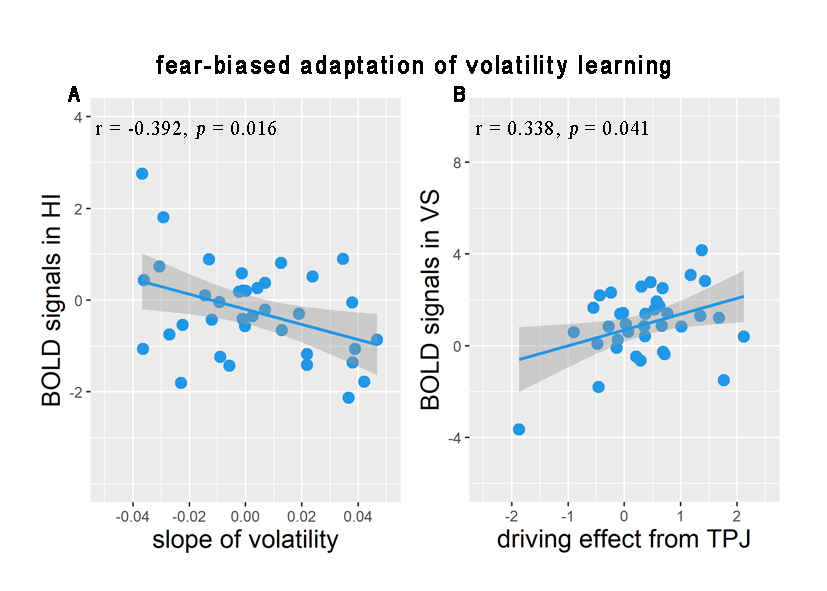

**Table S1.** Normative rating of fearful and neutral facial expressions from the Taiwanese Facial Expression Image Database (TFEID).

| Gender | Model | Category | Correction rate (%) | Intensity (0-8) |
| --- | --- | --- | --- | --- |
| Female | F05 | Neutral | 99.1 | 0.42 |
|  | F08 | Neutral | 100.0 | 0.41 |
|  | F04 | Fear | 86.0 | 4.85 |
|  | F18 | Fear | 73.9 | 5.12 |
| Male | M04 | Neutral | 97.3 | 0.46 |
|  | M12 | Neutral | 98.2 | 0.27 |
|  | M05 | Fear | 81.0 | 5.19 |
|  | M08 | Fear | 75.9 | 5.83 |

**Table S2.** Descriptive data of post-ratings and task performance.

|  | exp1 (n = 21) | | exp2 (n = 40) | |
| --- | --- | --- | --- | --- |
|  | Mean (SD) | Range [min,max] | Mean (SD) | Range [min,max] |
| Identification rating (fear) | 6.40 (1.15) | [4,8] | 6.16 (1.65) | [2,8] |
| Identification rating (neutral) | 6.64 (1.17) | [3,8] | 6.80 (1.06) | [4,8] |
| Number of missing trials | 1.76 (1.84) | [0,6] | 2.33 (3.52) | [0,14] |
| Ratio (%) of short response time (<200 ms) | 0. 2 (0. 6) | [0, 2.5] | 0.07 (0.28) | [0,1.67] |

**Table S3**. Behavioral and BOLD responses in each experimental condition.

| Variables | fear & freq | fear && infreq | neut & freq | neut & infreq |
| --- | --- | --- | --- | --- |
| learning rate (exp1) | 0.546 (0.239) | 0.654 (0.050) | 0.695 (0.104) | 0.541 (0.223) |
| learning rate (exp2) | 0.488 (0.242) | 0.562 (0.233) | 0.644 (0.198) | 0.463 (0.226) |
| volatility activation in VS | 0.313 (1.578) | -0.117 (1.342) | -0.201 (1.210) | 0.324 (1.015) |
| volatility activation in dACC | 0.408 (0.393) | -0.158 (2.214) | -0.915 (1.860) | 0.217 (2.331) |
| FC between dACC and AG | 0.477 (2.183) | -0.385 (1.622) | -0.490 (1.572) | 0.819 (1.600) |
| driving effect on AG | 0.093 (0.552) | 0.061 (0.488) | -0.286 (0.445) | 0.061 (0.488) |

Note: Descriptive data are presented as mean (SD).
